# A cell-free antibody engineering platform rapidly generates SARS-CoV-2 neutralizing antibodies

**DOI:** 10.1101/2020.10.29.361287

**Authors:** Xun Chen, Matteo Gentili, Nir Hacohen, Aviv Regev

**Author notes:** To whom correspondence should be addressed (X.C.), (A.R.).

## Abstract

Antibody engineering technologies face increasing demands for speed, reliability and scale. We developed CeVICA, a cell-free antibody engineering platform that integrates a novel generation method and design for camelid heavy-chain antibody VHH domain-based synthetic libraries, optimized *in vitro* selection based on ribosome display and a computational pipeline for binder prediction based on CDR-directed clustering. We applied CeVICA to engineer antibodies against the Receptor Binding Domain (RBD) of the SARS-CoV-2 spike proteins and identified >800 predicted binder families. Among 14 experimentally-tested binders, 6 showed inhibition of pseudotyped virus infection. Antibody affinity maturation further increased binding affinity and potency of inhibition. Additionally, the unique capability of CeVICA for efficient and comprehensive binder prediction allowed retrospective validation of the fitness of our synthetic VHH library design and revealed direction for future refinement. CeVICA offers an integrated solution to rapid generation of divergent synthetic antibodies with tunable affinities *in vitro* and may serve as the basis for automated and highly parallel antibody generation.

---

Antibodies and their functional domains play key roles in research, diagnostics and therapeutics. Antibodies are traditionally made by immunizing animals with the desired target as antigen, but such methods are time consuming, their outcome is often unpredictable, and their use is increasingly restricted in the European Union ^1^. Alternatively, antibodies can be generated and selected *in vitro*, where libraries of antibody-encoding DNA, either fully synthetic or derived from animals, are displayed *in vitro* followed by selection and recovery of those binding the intended target ^2,3^. However, broad application of such *in vitro* methods remains a challenge, possibly due to throughput limitations and concerns over functional fitness and *in vivo* tolerance of antibodies generated *in vitro* ^4^. Advances in antibody library design and construction, *in vitro* display and selection methods, post-selection binder identification and maturation will all help increase the utility of *in vitro* antibody generation ^2^.

For typical antibodies, antigen binding is co-determined by the variable domains of both its heavy chain (VH) and light chain (VL/VK), but camelids produce unconventional heavy-chain-only antibodies that bind to antigens solely based on the variable domain of their heavy chain, the VHH domain (also known as nanobodies). VHHs are increasingly used as functional antibody domains because of their small size (~14 kD) ^5^ and high stability (*T_m_* up to 90°C) ^6^. VHH libraries have been successfully screened for binders by phage and yeast display ^7–9^. However, the screen diversity of such cell-based systems is often limited by limited efficiency of DNA library delivery into cells (typically <10^10^). Conversely, cell-free approaches, such as ribosome display ^10^, are not limited by transfection efficiency and cell culture constraints. Despite the advantage, ribosome display remains underutilized compared to cell-based display systems ^2^ and recent efforts to build *in vitro* system based on ribosome display alone produced inconsistent results ^11^, suggesting that further optimization is required.

To leverage the advantages of cell-free display, we developed CeVICA (**<u>Ce</u>**ll-free **<u>V</u>**HH **<u>I</u>**dentification using **<u>C</u>**lustering **<u>A</u>**nalysis) (**Fig. 1**), an integrated platform for *in vitro* VHH domain antibody engineering, distinct from previous systems ^11–13^, that combines a novel design and generation method for CDR-randomized VHH libraries, optimized ribosome display and selection cycle with built-in background reduction, and a computational approach to perform global binder prediction from post-selection libraries. CeVICA first takes as input a linear DNA library, in which each sequence is unique and encodes for an artificial VHH with three fully-randomized CDRs, and where the 5’ and 3’ ends of the DNA molecules contain elements required for downstream *in vitro* ribosome display (**Fig. 1a, Materials and Methods**). Next, CeVICA uses ribosome display to link genotype (RNAs transcribed from DNA input library that are stop codon free, and stall ribosome at the end of the transcript) and phenotype (folded VHH protein tethered to ribosomes due to the lack of stop codon in the RNA) (**Fig. 1b, Materials and Methods**). In each selection cycle (**Fig. 1c, Materials and Methods**), the displaying ribosomes bind to an immobilized target, followed by RT-PCR of the RNA attached to the bound ribosomes, which leads to double stranded cDNA, which is then *in vitro* transcribed/translated in a new round of ribosome display. The double stranded DNA in any chosen round is sequenced to obtain full-length VHH sequences (**Fig. 1d, Materials and Methods**). CeVICA then groups the sequences into clusters based on similarity of their CDR sequences, such that each cluster represents a unique binding family (**Fig. 1e, Materials and Methods**). Finally, one representative sequence from each cluster is synthesized and characterized for specific downstream applications (**Fig. 1f, Materials and Methods**). The combination of linear DNA libraries (**Fig. 1a**), ribosome display (**Fig. 1b**) and selection cycles (**Fig. 1c**) allow display of libraries with much larger diversity (>10^10^) than methods depending on cells ^14^ at similar experimental scale. As selection increases the representation of sequences encoding binders, each binder sequence leads to a cluster of sequences in the output library. Clustering following high throughput sequencing identifies them more efficiently than methods that rely on the analysis of individual colonies or sequences ^7,8^, promising a more comprehensive view of the landscape of binder potential, with minimal time and resources.

**Fig. 1.**
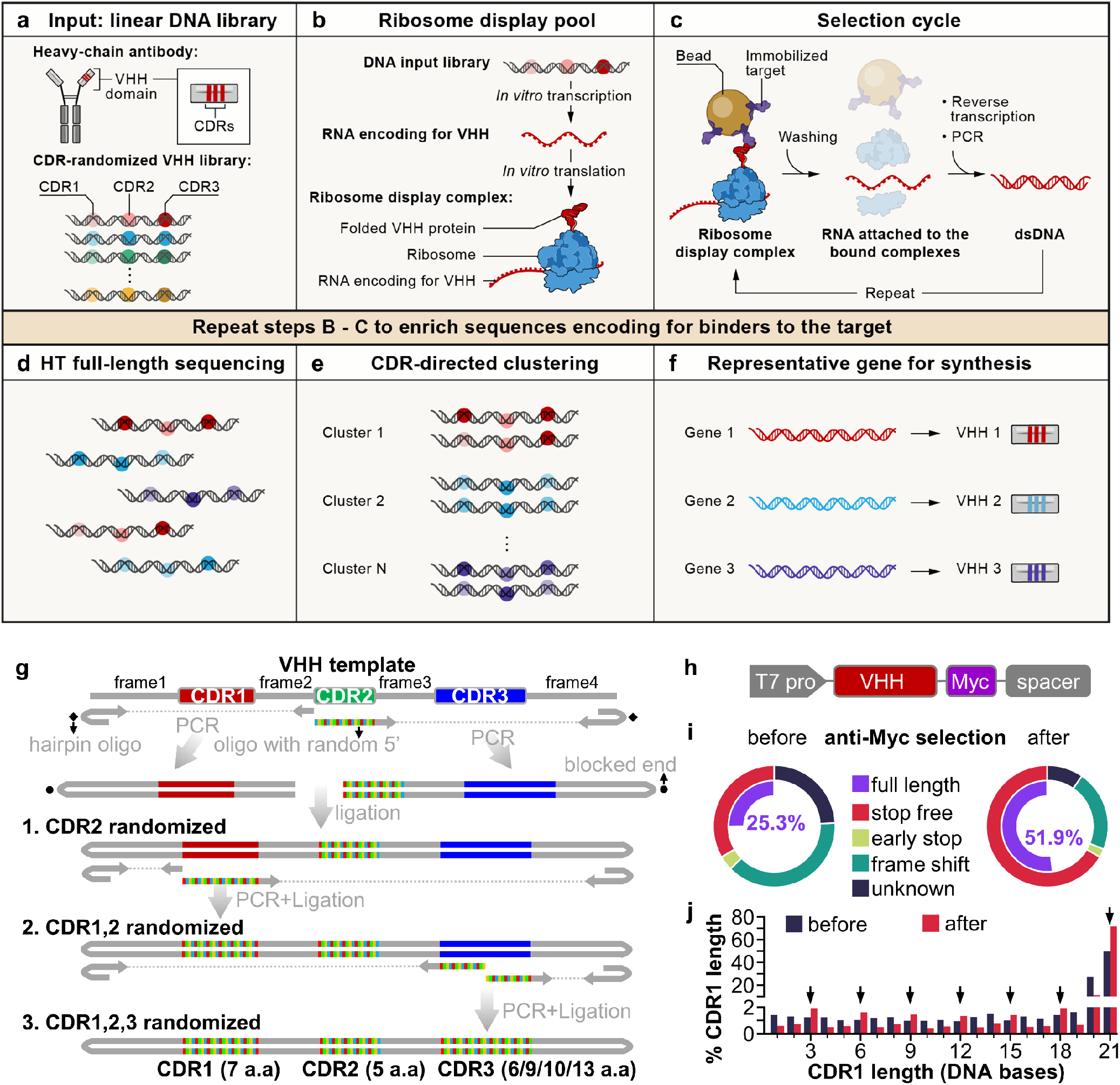
A cell-free antibody engineering platform for rapid isolation of antibodies from large synthetic libraries. (**a**) The workflow takes linear DNA library as input. (**b**) Ribosome display links genotype (RNAs transcribed from DNA input library that are stop codon free, and stall ribosome at the end of the transcript) and phenotype (folded VHH protein tethered to ribosomes due to the lack of stop codon in the RNA). (**c**) Selection cycle that enriches DNA encoding for VHHs that binds immobilized targets. (**d**) High throughput sequencing of full-length VHHs. (**e**) Sequences are grouped into clusters based on similarity of their CDRs, each cluster is distinct and represent a unique binding family. (**f**) The system outputs one representative sequence from each cluster to be synthesized and characterized for specific downstream applications. (**g**) Workflow for generating VHH library. VHH CDR randomization was introduced by PCR using a hairpin oligo (blocks DNA end from ligation) and an oligo with random 5’ sequence, followed by orientation-controlled ligation. Three successive PCR plus ligation cycles randomizes all three CDRs. (**h**) The final DNA library sequence structure. (**i**) One round of ribosome display and anti-Myc selection was performed after randomization of CDR1 and CDR2. The pie chart shows percentage of indicated sequence categories before and after anti-Myc selection. (**j**) Length distribution of DNA region encoding CDR1 of the VHH library before and after anti-Myc selection. Arrows indicate all correct-frame lengths showing increased percentage after anti-Myc selection.

We made VHH libraries containing highly random CDRs, based on analysis of natural VHH sequences and using a three-stage PCR and ligation process (**Fig. 1g**). First, to guide our VHH library sequence design, we analyzed the sequence characteristics of 298 unique camelid VHHs (representing natural VHHs) from the Protein Data Bank (PDB) (**table S1**), highlighting three CDR regions, CDR1-3 ^5^, separated by four regions of low diversity, frame1-4 (**Fig. S1a**). The four frames share high homology with human IGHV3-23 or IGHJ4 (**Fig. S2a,b**), and most of the remaining non-identical residues are present in other human IGHV genes (**Fig. S2c**). We used consensus sequences extracted from this profile to design VHH DNA templates encoding the four frames (**Fig. 1g**), and included additional frames to the final mixture of frame templates (**Materials and Methods**), based on well-characterized VHHs ^6,15^. The mixture of VHH frames serves as a template in PCR reactions, where DNA oligonucleotides with a 5’ NNB sequence were used to introduce randomization in CDRs, while hairpin DNA oligonucleotides were used to block ligation of one end of the PCR product (**Fig. 1g** and **Fig. S3**, **Materials and Methods**). We introduced 7 random amino acids for CDR1, 5 for CDR2, and 6, 9, 10 or 13 for CDR3 to match the most commonly observed CDR lengths in natural VHHs. CDR3s longer than 13 amino acids only account for a minority of natural VHHs (36%, **Fig. S1a**, **table S2**) and were not included in our VHH library. CDRs randomized in earlier stages are subject to duplication in later stages that reduces their diversity. We thus chose to randomize CDR2 first, followed by CDR1, and then CDR3, imposing a diversity hierarchy of CDR3>CDR1>CDR2, because this is the overall ranking of diversity we observed in CDRs in natural VHHs (**Fig. S1a,c**). The sequence profile of the resulting randomized VHH library met our design objectives, and largely mirrored the sequence features of natural VHHs (**Fig. S1** and **table S2**). Finally, the VHH DNA library contains an upstream T7 promoter to allow transcription of VHH RNA, a 3xMyc tag, and a spacer downstream of the VHH coding region that stalls peptide release, to enable ribosome display (**Fig. 1h**).

To test the performance of our library in ribosome display, and to reduce unproductive sequences, such as VHHs that contain frame shifts or early stops, we ribosome displayed a library only with randomized CDR1 and CDR2 and performed one round of anti-Myc selection. Functional VHH sequences will express Myc tag at the C-terminal of VHH and are expected to be enriched after anti-Myc selection. Indeed, there was a large decrease of unproductive sequences and an increase of full-length VHHs (from 25.3% to 51.9%) after anti-Myc enrichment (**Fig. 1i**). At the DNA level, there was an increase of all in-frame CDR1 DNA lengths and decrease of frame-shift lengths (**Fig. 1j,**arrows). We used the resulting full-length enriched CDR1 and 2 randomized library as PCR template for randomization of CDR3. The final library with all three CDRs randomized (hereafter, “the input library”) contained 27.5% full-length sequences, and 3.68×10^11^ full-length diversity per μg of library DNA.

We performed *in vitro* selection from the input library for sequences that encode binders to two target proteins: EGFP and the receptor binding domain (RBD) of the spike protein of SARS-CoV-2 ^16^ (**Fig. 2**). We fused each of the two proteins with a 3xFlag tag and immobilized them on beads coated with protein G and anti-Flag antibody (**Fig. 2a**). For each screen, we used input library DNA corresponding to ~1×10^11^ full-length diversity, and performed 3 rounds of selection. After round 3, RNA yield markedly increased in both screens (**Fig. S4a**) and the recovered sequences were primarily composed of *E. coli* ribosomal RNAs and VHH library RNA (*e.g.*, **Fig. S4b**). Comparing the input and output library sequences shows a marked increase in the proportion of stop-free VHH sequences after 3 rounds of selection (**Fig. 2c**), fitting our expectation that successful binding to targets depends on intact VHH structure.

**Fig. 2.**
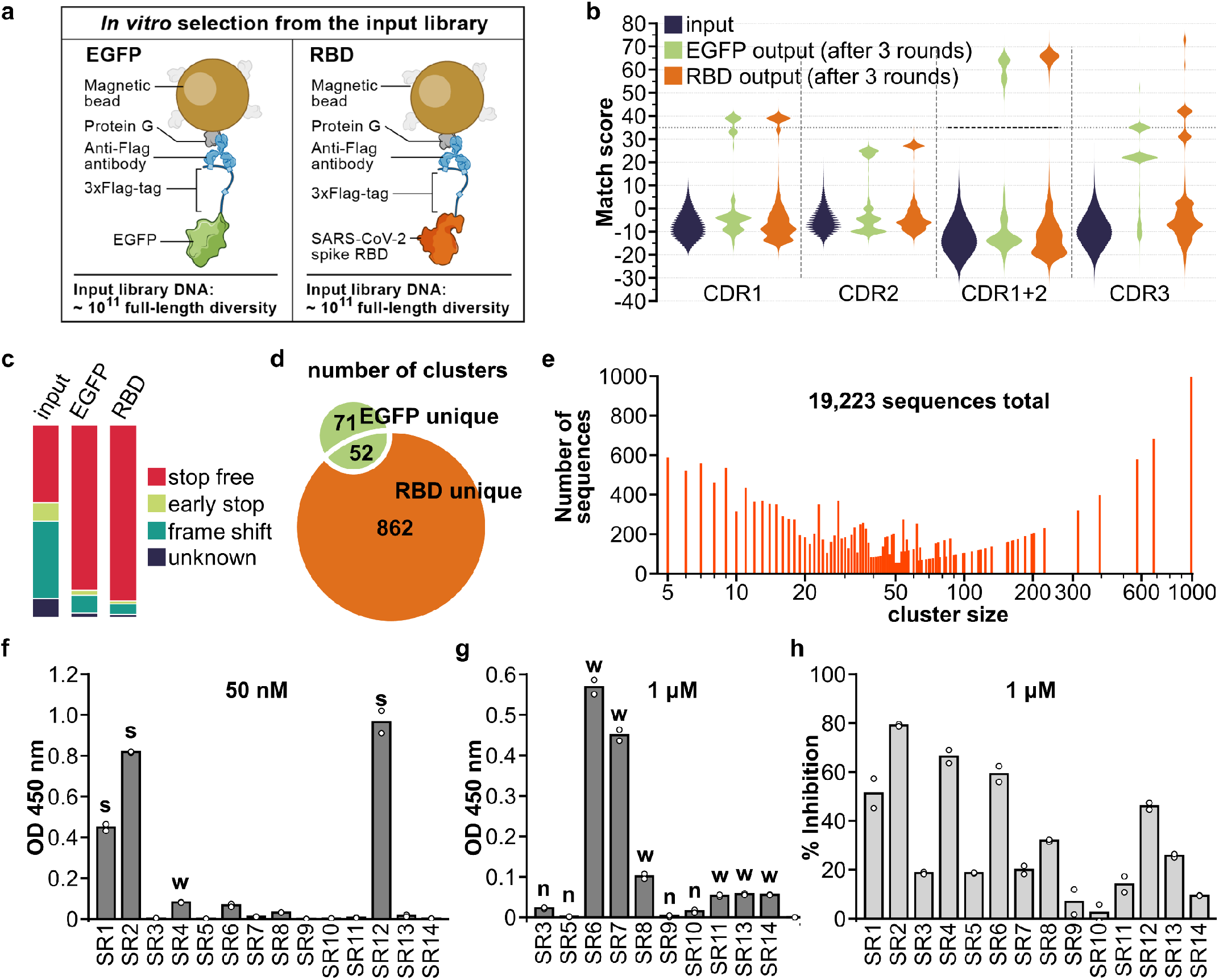
Isolation and characterization of synthetic VHHs that binds SARS-CoV-2 spike RBD. (**a**) Immobilization strategy for the target proteins: 3xFlag-tagged EGFP or RBD. (**b**) Pair-wise CDR match score (based on BLOSUM62 matrix) were calculated for 2000 randomly selected sequences from input library and output libraries after 3 rounds of selection. High match score populations appeared in the output libraries. Combining CDR1 and 2 match scores further separated high and low score population and a match score of 35 (black dashed line) was chosen as cut-off for downstream clustering analysis. (**c**) Percentage of indicated sequence categories in the input library and output libraries (EGFP, RBD). (**d**) Number of unique and shared clusters identified in EGFP and RBD output libraries. (**e**) Number of sequences for each size of RBD unique clusters. (**f**) ELISA assay revealed 3 strong binders (“s”) to RBD, 7 weak binders (“w”) and (**g**) 4 non-binders (“n”) among the 14 VHHs chosen for characterization. (**h**) SARS-CoV-2 S pseudotyped lentivirus neutralization assay showed 6 VHHs inhibiting infection >30% at 1μM on HEK293T expressing ACE2 and TMPRSS2. Data shown are two technical replicates, bars indicate the average of data, circles indicate values of each replicate.

We identified target specific binders by clustering CDR sequences enriched after selection into families. First, we examined the distribution of the sequence match scores (**Materials and Methods**) between randomly selected pairs of sequences within a CDR in a library, and compared these distributions for each CDR between the input and output libraries (**Fig. 2b, Materials and Methods**). In the pre-selection input libraries, the mean match score is low and the distribution is unimodal, as expected given the randomization; whereas after selection, there is a multi-modal distribution, with one low mode (similar to input) and at least one high mode (**Fig. 2b**), which is further distinguished when combining the CDR1 and CDR2 match scores (**Fig. 2b**). This high mode should reflect binders enriched by the selection rounds. Notably, sequences with a high match score in one CDR are more likely to have a higher match score in other CDRs (**Fig S4c-f**). We clustered the likely binder sequences exceeding a combined match score threshold (**Fig. 2b**, dashed horizontal line), yielding 862 unique clusters for RBD and 71 for EGFP, with 52 clusters shared by the two targets (**Fig. 2d, table S4 and 5**). The shared clusters likely target the shared components (protein G, anti-Flag antibody) present on the solid support surfaces, and thus represent background binders. Notably, RBD unique clusters span a wide range of cluster sizes (**Fig. 2e**).

Focusing on RBD binders, we chose one representative VHH gene from each of the14 top-ranking RBD unique clusters and validated it for spike RBD binding and SARS-CoV-2 pseudovirus neutralization (**Fig. 2f-h**, **Materials and Methods**). RBD binding ELISA assays of the 14 tested VHHs (SR1-14) showed 3 strong binders (SR1,2,12), 7 weak binders (SR4,6,7,8,11,13,14) and 4 non-binders (**Fig. 2f,g**). SARS-CoV-2 S pseudotyped lentivirus neutralization assays revealed 6 VHHs inhibiting infection above 30% at 1 μM (**Fig. 2h**), which included the 3 strong binders and three of the weak binders (SR4,6,8).

We next compared input, output and natural CDR sequence distributions to assess whether starting with a fully random CDR amino acid profile may be generally detrimental to the fitness of binders, and whether selection mimics a natural amino acid distribution. In natural VHHs, CDR1 and CDR2 are less diverse than CDR3 with an amino acid profile that favors certain residues (**Fig. S1a,c**). Previous synthetic VHH library designs sought to recapitulate the CDR1 and CDR2 amino acid preferences of natural VHHs ^8,11,13^, whereas we used fully-randomized NNB codons to encode all CDR positions. In principle, such a design might be less ideal if the natural CDR1 and CDR2 amino acid profile is required for functional VHHs. To determine whether our fully random CDR amino acid profile is detrimental to the fitness of binders, we compared the CDR amino acid profile of 932 representative sequences across all unique clusters from both the EGFP and RBD output libraries (“output binders”) (**Fig. S5**) to the sequence profiles of either the input library or natural VHHs (**Fig. S1a,b**). We reasoned that if the amino acid profile in the input library leads to a distribution of proteins that are less fit in binding, the binder selection process should shift this distribution to a more fit profile in the output library, such that there is a low correlation between the amino acid profiles of the input library and output binders. Surprisingly, there was an overall smaller shift in CDR1 and CDR2 compared to CDR3, as indicated by higher *r*^2^ values (**Fig. S6a-c, mean***r*^2^ = 0.45, 0.51, and 0.36 respectively), and lower similarity distances (as the RMSE relative to y = x line, **Materials and Methods**, **Fig. S6d,e**, RMSE = 2.96, 2.40 and 3.51 respectively), implying that a fully random profile at CDR1 and CDR2 may not have had a substantial binding fitness cost at most positions, whereas CDR3 not only shifted away from the input profile, it was even further shifted from the natural profile (**Fig. S6d,e**). Moreover, correlation of amino acid profiles between output binders and natural VHHs are significantly less than between output binders and input library at most CDR positions (**Fig. S6**). A few positions (CDR1 position 7 and CDR3 position 1-3) had much lower input-output binders *r*^2^ than others. This suggests that these positions may benefit from specifically-designed amino acid profiles (to adjust off diagonal amino acids percentages (**Fig. S6b**) accordingly), even though their input distributions were not particularly distinct from the native sequence distribution compared to other positions (**Fig. S6a,d**). Thus, the output binder CDR profile is predominantly influenced by the input library rather than by selection towards a natural VHH profile, a natural VHH CDR amino acid profile is not required for VHH binding properties, and a fully random CDR design offers high diversity without a major binding fitness cost (although may have other fitness drawbacks *in vivo*).

To perform affinity maturation, a critical stage in antibody development in animals, we designed and performed an affinity maturation strategy based on CeVICA to increase the affinity of RBD binding VHHs (**Fig. 3a, Materials and Methods**). We used error-prone PCR to introduce random mutations across the full-length sequence of six selected VHHs (SR1,2,4,6,8,12) and generated the mutagenized library. A library size of 4.18×10^10^ diversity (sufficient to contain the full diversity of VHHs with three mutations per sequence) was used as input and three rounds of stringent selection were performed. We sequenced the libraries pre- and post-affinity maturation, and observed about 3 mutations in the pre-library and about 2 mutations in the post-library per sequence (**Fig. 3a**). We calculated their position-wise amino acid profiles, and determined, for each VHH, the change in each amino acid proportion at each position, generating a percent point change table. We defined putative beneficial mutations as those with a percent point increase above a set threshold (**Fig. 3b, Materials and Methods** and **table S6**), highlighting between 8 to 25 putative beneficial mutations for each of the selected VHHs. Finally, we assembled a list of identified putative beneficial mutations for each VHH and incorporated different combinations of them into each VHH parental sequence to generate multiple mutated variants of each VHH for final assessment (**table S7**).

**Fig. 3.**
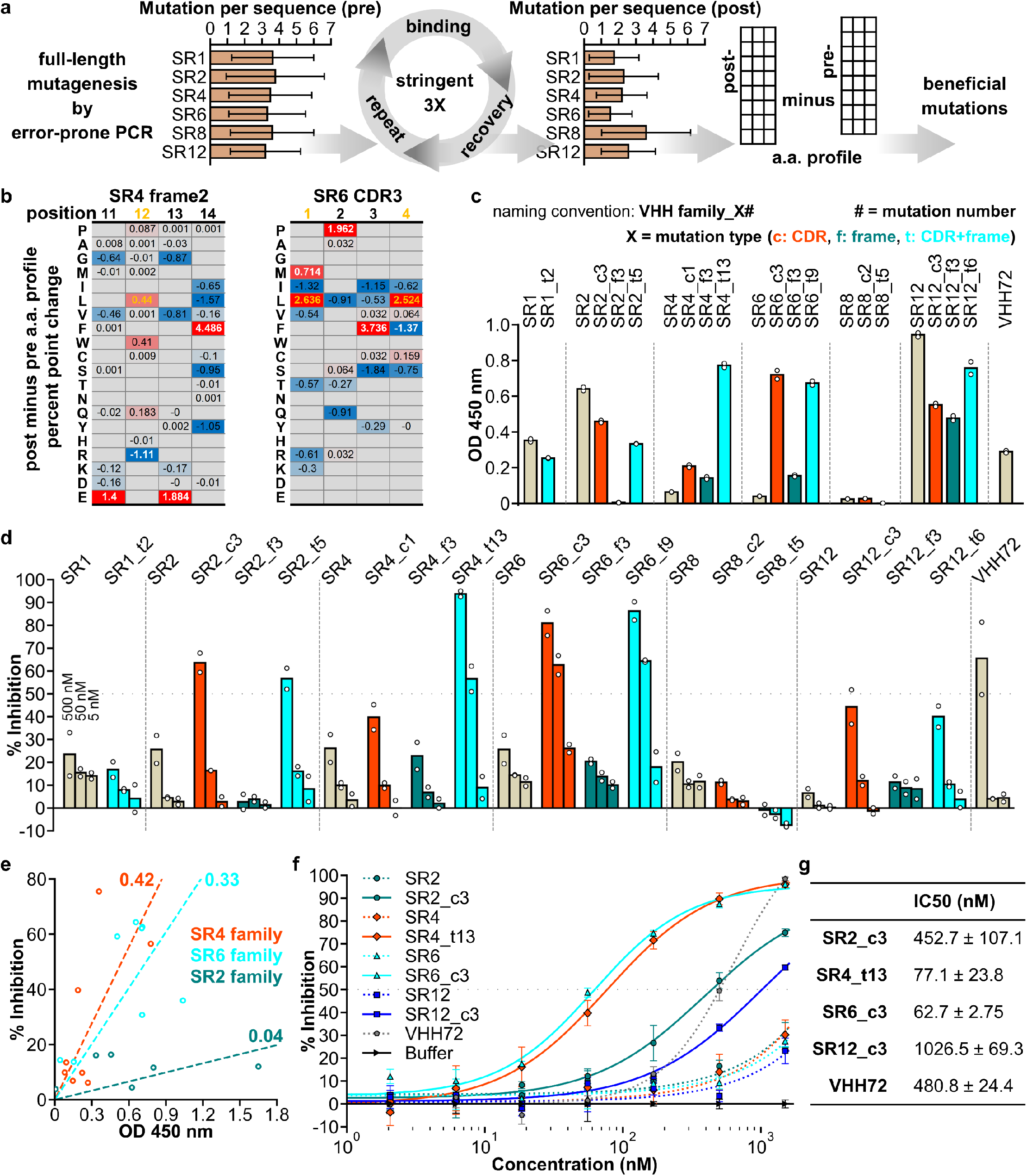
An affinity maturation strategy enhances binding and neutralization properties of synthetic VHHs. (**a**) Affinity maturation workflow. (**b**) Two representative sections of position-wise post-minus pre-affinity maturation amino acid percent point change profile. White values indicate the original amino acid, yellow values indicate the beneficial mutation. Empty positions indicate amino acids not detected in either the pre-or post-selection libraries. (**c**) ELISA assay of VHH variants. (**d**) SARS-CoV-2 S pseudotyped lentivirus neutralization assay of VHHs on HEK293T expressing ACE2 and TMPRSS2. For (c) and (d), data shown are two technical replicates, bars indicate the average of data, circles indicate values of each replicate. (**e**) Scatter plot of ELISA assay absorbance versus pseudotyped lentivirus neutralization as percent infection inhibited. VHH concentration for both assays were 50 nM. Values are average of two technical replicates. Numbers on linear fitting lines were *r*^2^ value for data within each family. (**f**) Dose-response curve for neutralization of pseudotyped lentiviral infection by VHHs. Markers are average of three technical replicates, error bars are standard deviation. (**g**) IC50 calculated from data in (f), presented as mean ± standard deviation.

Variants in the SR4 and SR6 families had both increased binding and neutralization, while the SR2 and SR12 family variants had only increased neutralization but not binding, based on an ELISA binding assay and a pseudotyped virus neutralization assay (**Fig. 3c,d**). Multiple VHH variants outperformed VHH72, a previously described VHH antibody that neutralizes SARS-CoV-2 pseudoviruses (Wrapp et al., 2020), in binding (*e.g.*, SR12_c3), neutralization (*e.g.*, SR4_t6), or both (*e.g.*, SR6_c3) (**Fig. 3c,d** and **table S8**). Neutralization and binding performance were poorly correlated across variants (*r*^2^ = 0.07), as previously reported ^17^. However, when considering each VHH family separately, trends were stronger, and neutralization and affinity were more highly correlated for SR4 and SR6 VHHs (**Fig. 3e**). This may be because variants within the same family share the same binding site and orientation. One intriguing hypothesis is that the slope of each VHH family’s linear trend reflects the sensitivity of the virus to the blocking of the family’s binding site. A dose response curve of selected VHHs showed SR6_c3 as the most potent neutralizer (**Fig. 3f**) with an IC50 of 62.7 nM (**Fig. 3g**), comparable to the Fab domains of potent SARS-CoV-2 neutralizing antibodies identified from human patients ^18^. Importantly, the original SR6 cluster contained only 679 sequences, representing 0.67% of the 101,674 sequenced from the initial selection output, highlighting the power of CeVICA in rapidly identifying high performance antibodies among a vast number of potential candidates.

Finally, we examined the potential impact that our VHH sequences may have on immunogenicity in humans, as a major concern related to the therapeutic use of VHH antibodies is the possibility that, as camelid proteins, they would elicit an immune response. In particular, VHH hallmark residues in frame2 constitute a major difference between camelid VHHs and human VHs (**Fig. S2**). We used our affinity maturation data to identify potential conversion options for these VHH hallmark residues. In three of the four VHH hallmark residues there were VHHs where the residues were converted to the corresponding human residue as a result of affinity maturation (**Fig. S7**, arrows). These data imply that at least some of the VHH hallmark residues can be converted to human residues without loss of binding fitness. Such conversions may serve as frame features of future VHH library designs and improve tolerance of *in vitro* engineered VHHs by humans. Overall, the extension of CeVICA for affinity maturation offers a strategy for improving antibody function and additional iterations of the affinity maturation process may provide further enhancement of antibody properties.

In conclusion, CeVICA is a new system for synthetic VHH based antibody library design, *in vitro* selection optimization, post-selection screening, and affinity maturation. Using CeVICA, we generated a large collection of antibodies that can bind the RBD domain of the SARS-CoV-2 spike protein and can neutralize pseudotyped virus infection, thus providing an important resource. Given its seamlessly integrated procedure, CeVICA is amenable to automation and could provide an important tool for antibody generation in a rapid, reliable and scalable manner. CeVICA further provides a technology framework for incorporation of future refinements that could overcome limitations of *in vivo* fitness of *in vitro* generated antibodies and overall efficiency.

## Materials and Methods

### Constructs

DNA encoding VHHs were obtained by gene synthesis (IDT) and cloned into pET vector in frame with a C-terminal 6XHis tag by Gibson assembly (NEBuilder® HiFi DNA Assembly Master Mix, New England Biolabs). DNA encoding SARS-CoV-2 S RBD (S a.a. 319-541) were obtained by gene synthesis and cloned into pcDNA3 with an N-terminal SARS-CoV-2 S signal peptide (S a.a. 1-16) and a C-terminal 3xFlag tag by Gibson assembly. EGFP was cloned into pcDNA3 with a C-terminal 3xFlag tag by Gibson assembly. SARS-CoV-2 S was amplified by PCR (Q5 High-Fidelity 2X Master Mix, New England Biolabs) from pUC57-nCoV-S (kind gift from Jonathan Abraham lab). SARS-CoV-2 S was deleted of the 27 a.a. at the C-terminal and fused to the NRVRQGYS sequence of HIV-1, a strategy previously described for retroviruses pseudotyped with SARS-CoV S ^19^. Truncated SARS-CoV-2 S fused to gp41 was cloned into pCMV by Gibson assembly to obtain pCMV-SARS2ΔC-gp41. psPAX2 and pCMV-VSV-G were previously described ^20^. pTRIP-SFFV-EGFP-NLS was previously described ^21^ (a gift from Nicolas Manel; Addgene plasmid # 86677; http://n2t.net/addgene:86677; RRID:Addgene_86677). cDNA for human TMPRSS2 and Hygromycin resistance gene was obtained by synthesis (IDT). pTRIP-SFFV-Hygro-2A-TMPRSS2 was obtained by Gibson assembly.

### Cell culture

HEK293T cells were cultured in DMEM, 10% FBS (ThermoFisher Scientific), PenStrep (ThermoFisher Scientific). HEK293T ACE2 were a kind gift of Michael Farzan. HEK293T ACE2 cells were transduced with pTRIP-SFFV-Hygro-TMPRSS2 to obtain HEK293T ACE2/TMPRSS2 cells. The transduced cells were selected with 320 μg/ml of Hygromycin (Invivogen) and used as a target in SARS-CoV-2 S pseudotyped lentivirus neutralization assays. Transient transfection of HEK293T cells was performed using TransIT®-293 Transfection Reagent (Mirus Bio, MIR 2700).

### Amino acid profile construction and analysis of natural VHHs

VHH protein sequences were downloaded from the Protein Data Bank (only entries deposited prior to Sep 2^nd^, 2020 were included; **table S1**). VHHs were separated into CDRs and frames (segments) by finding regions of continuous sequence in each VHH that best matched to the following standard frame sequences:

frame1 standard: EVQLVESGGGLVQAGDSLRLSCTASG,
frame2 standard: MGWFRQAPGKEREFVAAIS,
frame3 standard: AFYADSVRGRFSISADSAKNTVYLQMNSLKPEDTAVYYCAA,
frame4 standard: DYWGQGTQVTVSS,

Each matched region is the corresponding frame of the VHH, the region between frame1 and frame2 is CDR1, the region between frame2 and frame3 is CDR2, the region between frame3 and frame4 is CDR3 (**Fig. 1g**). Only VHH sequences with at least one unique CDR were selected to represent natural VHHs and used for constructing amino acid profile (a.a. profile). 298 sequences fit this selection criteria (**table S1**). The amino acid (a.a.) profile at each position within each segment was calculated by finding the percentage of each of the 20 universal proteinogenic amino acid at that position among all selected VHHs, all frame lengths were set to the same length as frame standards. CDR lengths were manually set to accommodate different CDR lengths, CDR1 and CDR2 lengths was set to 10, CDR3 length was set to 30. VHHs with CDR lengths shorter than the corresponding set length had their CDR filled from the C-terminal end with empty position holders up to the set length. Numbers in amino acid profile table are the percentage of each amino acid.

### VHH library construction

VHH libraries were constructed by ligation of PCR products in three stages, with each stage randomizing one of the three CDRs. Primers used and PCR cycling conditions for each primer pair are listed in **table S3**. At each stage, PCR was performed using a high-fidelity DNA polymerase without strand displacement activity, using Phusion DNA polymerase (New England Biolabs, M0530L). Importantly, 65°C was used as the elongation temperature to avoid hairpin opening during DNA elongation. PCR products with correct size were purified by DNA agarose gel extraction. Ligation and phosphorylation of PCR products were performed simultaneously using T4 DNA ligase (New England Biolabs, M0202L) and T4 Polynucleotide Kinase (New England Biolabs, M0201L). Ligation products with the correct size were purified by DNA agarose gel extraction using NucleoSpin Gel and PCR Clean-Up Kit (Takara, 740609.250, this kit was used for all DNA agarose gel extraction steps in this study). Purified ligation products were quantified with Qubit 1X dsDNA HS Assay Kit (ThermoFisher Scientific, Q33230, this kit was used for all Qubit measurements in this study) using Qubit 3 Fluorometer.

CDR2 was randomized in stage one, PCR templates at this stage were equal molar mixtures of plasmids carrying DNA encoding frames, including three frame1 versions, one frame2, three frame3 versions and one frame4. The three versions of frame1 and frame3 were derived from consensus sequence extracted from natural VHH a.a. profile, the A3 VHH ^6^ and a GFP binding VHH ^15^. Amino acid sequences of the frames are shown in **fig. S1**.

CDR1 was randomized in stage two, 200 ng of ligation product from the first stage were digested by Not I-HF (New England Biolabs, R3189S) and heat denatured, the entire digestion product was used as template for PCR in stage two. Ligation product of stage two was subject to one round of ribosome display and anti-Myc selection (below), the entire recovered RNA was reverse transcribed and PCR amplified and purified.

270 ng of this RT-PCR product was used as template for PCR in stage three to randomize CDR3. Ligation product of stage three was purified by DNA agarose gel extraction. The purified ligation product was then digested by DraI (New England Biolabs, R0129S) and a fragment of ~680 bp in size was purified by DNA agarose gel extraction to get the final VHH library, referred to as the input library.

### High throughput full-length sequencing of VHH library

Sequencing libraries from VHH DNA libraries were prepared by two PCR steps using primers and PCR cycling conditions listed in **table S3**. Equal mixtures of Phusion DNA polymerase (New England Biolabs, M0530L) and Deep Vent DNA polymerase (New England Biolabs, M0258L) were used for both PCRs to ensure efficient amplification. PCR cycle number was chosen to avoid over-amplification and typically falls between 5 to 15.

In the first PCR, Illumina universal library amplification primer binding sequence and a stretch of variable lengths of random nucleotides were introduced to the 5’ end of library DNA. And similarly, Illumina universal library amplification primer binding sequence and a stretch of variable lengths of index sequence are introduced to the 3’ end of library DNA. Eight different lengths were used for both random nucleotides and index to create staggered VHH sequences in the sequencing library, this arrangement is required for high quality sequencing of single amplicon libraries on an Illumina Miseq instrument. The product of the first PCR was purified by column clean-up using NucleoSpin Gel and PCR Clean-Up Kit and the entire sample was used as template for the second PCR.

In the second PCR, Illumina universal library amplification primers were used to generate sequencing library. Sequencing libraries were purified by DNA agarose gel extraction, quantified using Qubit 3 Fluorometer, and sequenced on an Illumina Miseq instrument using MiSeq Reagent Nano Kit v2 (500-cycles) (Illumina, MS-103-1003), no PhiX control library spike-in was used. Sequencing run setup was: paired end 2X258 with no index read. Index in the library was designed as inline index, so a separate index read was not required.

### Ribosome display

VHH DNA library containing a specified amount of diversity was first amplified using a DNA recovery primer pair listed in **table S3**. Equal mixtures of Phusion DNA polymerase (New England Biolabs, M0530L) and Deep Vent DNA polymerase (New England Biolabs, M0258L) were used for the PCR. PCR cycle number was chosen to avoid over-amplification and typically falls between 5 and 15. In a standard preparation, 200-500 ng of the purified PCR product was used as DNA template in 25 μl of coupled *in vitro* transcription and translation reaction using PURExpress In Vitro Protein Synthesis Kit (New England Biolabs, E6800L). The reaction was incubated at 37°C for 30 minutes, then placed on ice, and 200 μl ice cold stop buffer (10 mM HEPES pH 7.4, 150 mM KCl, 2.5 mM MgCl_2_, 0.4 μg/μl BSA (New England Biolabs, B9000S), 0.4 U/μl SUPERase•In (ThermoFisher Scientific, AM2696), 0.05% TritonX-100) was then added to stop the reaction. This stopped ribosome display solution was used for binding to immobilized protein targets during *in vitro* selection. The amount of DNA template, volume of coupled *in vitro* transcription and translation reaction, and volume of stop buffer were scaled proportionally when different volumes of stopped ribosome display solution was needed. 1 to 8X standard preparations were used for each selection cycle.

### *In vitro* selection

Target proteins were immobilized to magnetic beads by first coating protein G magnetic beads (ThermoFisher Scientific, 10004D) with anti-Flag antibody (Sigma-Aldrich, F1804), then incubating antibody-coated beads with cell lysate or cell media containing 3xFlag tagged target proteins at 4°C for 2 hours. For anti-Myc selection, magnetic beads were coated by anti-Myc antibody (ThermoFisher Scientific, 13-2500) only. The beads were washed three times with PBST (PBS, ThermoFisher Scientific, with 0.02% TritonX-100). Beads were then incubated with stopped ribosome display solution at 4°C for 1 hour, and then washed 4 times with wash buffer (10 mM HEPES pH 7.4, 150 mM KCl, 5 mM MgCl_2_, 0.4 μg/μl BSA (New England Biolabs, B9000S), 0.1U/μl SUPERase•In (ThermoFisher Scientific, AM2696), 0.05% TritonX-100). After washing, beads were resuspended in TRIzol Reagent (ThermoFisher Scientific, 15596026), and RNA was extracted from the beads, 25 μg of linear acrylamide (ThermoFisher Scientific, AM9520) were used as co-precipitant during RNA extraction. Reverse transcription of extracted RNA was performed using Maxima H Minus Reverse Transcriptase (ThermoFisher Scientific, EP0752). The reverse transcription reaction was purified using SPRIselect Reagent (Beckman Coulter, B23317) to obtain purified cDNA. Purified cDNA was amplified by PCR using equal mixtures of Phusion DNA polymerase and Deep Vent DNA polymerase. PCR cycle number (**table S3**) was chosen to avoid over-amplification and typically falls between 10 to 25. The PCR product was purified by DNA agarose gel extraction. The purified PCR product was used for library generation for high throughput full-length sequencing or as DNA template for ribosome display reaction (coupled in vitro transcription and translation) to perform additional rounds of in vitro selection.

### CDR-directed clustering analysis

Computational analysis for CDR-directed clustering was performed using custom python scripts. Paired end sequences were merged to form full-length VHH sequences. Merged VHH sequences were quality trimmed and translated into VHH protein sequence, which were separated into CDRs and frames (segments) as described in the ***Amino Acid Profile Construction*** section. Two VHHs were determined to have similar CDRs via the following steps. First, the ungapped sequence alignment score (match score) was calculated for each CDR of the two VHHs as the sum of BLOSUM62 ^22^ amino acid pair scores at each aligned position. (If two CDRs have different lengths, their sequence alignment score was set to −5 by default.) The alignment scores of any two pairs of CDRs were summed to yield three scores, and if at least one of the three was larger than 35 (**Fig. 2b**), the two VHHs were defined as having similar CDRs. Next, VHHs with similar CDRs were grouped by a two-step process. In the first step, we chose as VHH cluster-forming “seeds” those VHHs that were called as similar to at least 5 other VHHs (all remaining VHHs were not considered for clustering). In the second step, we iteratively selected a seed VHH with at least 5 other similar (>35 match score) seed VHHs, and grouped all of them into one cluster, removing them from the seed VHH pool, and iterated this procedure until no seed VHHs remained. For RBD, there were 83,433 seeds in the first step, and 83,392 were grouped in clusters in the second step. For EGFP, 71,210 of 71,220 seeds were grouped in clusters (**table S9**). This heuristic was fast in a standard computing environment with multiprocessing capabilities.

A representative sequence to illustrate each CDR in each cluster was chosen as the most frequent CDR sequence in the cluster (the chose representatives for CDR1,2, and 3 may not necessarily be from the same sequence, and are used only for illustrative purposes for each cluster as in **table S4** and **S5**; whole VHH sequences were used for gene synthesis and all downstream experiments). A consensus sequence was generated for each CDR, where each position in the CDR was represented by a 6 character string, such that the first and fourth character were the single letter code for the top and the second most abundant amino acid at the position, respectively, and the following two characters (second and third for the most abundant; fifth and sixth for the second most abundant), were their frequency, respectively (ranging from 00 for <34% to 99 for 100%). The consensus sequence for a CDR was recorded as a single “B00” when the standard deviation of the lengths of all CDRs was greater than 1. CDR scores were calculated by summing a score for each position in the CDR consensus sequence, with scores of 3, 2, 1 for positions where the most abundant amino acid had frequencies greater than 80%, 50%, or less, respectively, and a score of 0 for CDRs with a consensus sequence of a single “B00” (**table S4** and **table S5**). Representative whole VHH sequence for each cluster was selected as the one with the maximal sum of all CDR similarity score between each VHH and all other VHHs in the cluster.

### Protein expression and purification

Target proteins used for *in vitro* selection and ELISA were prepared by transiently transfecting HEK293T cells with plasmids carrying either spike RBD with C-terminal 3xFlag tag and N-terminal signal peptide of spike (RBD-3xFlag), or EGFP with C-terminal 3xFlag tag (EGFP-3xFlag). Cell culture media (for RBD-3xFlag) or lysate of cell pellet (for EGFP-3xFlag) were used for coating magnetic beads or plates. VHHs with C-terminal 6XHis tag (VHH-6XHis) were purified by expressing in *E. coli.*, followed by purification using HisPur Cobalt Resin (ThermoFisher Scientific, 89964). Briefly, VHH-6xHis plasmids were transformed into T7 Express *E. Coli.* (New England Biolabs, C2566I), single colonies were transferred into 10 ml LB media and grown at 37°C for 2-4 hours (until OD reached 0.5-1), the culture was chilled on ice, then IPTG was added to a final concentration of 10 μM. The culture was then incubated on an orbital shaker at room temperature (RT) for 16 hours. Bacterial cells were pelleted by centrifugation and lysed in B-PER Bacterial Protein Extraction Reagent (ThermoFisher Scientific, 78248) supplemented with rLysozyme (Sigma-Aldrich, 71110), DNase I (New England Biolabs, M0303S), 2.5 mM MgCl_2_ and 0.5 mM CaCl_2_. Bacterial lysates were cleared by centrifugation and mixed with wash buffer (50 mM Sodium Phosphate pH 7.4, 300 mM Sodium Chloride, 10 mM imidazole) at 1:1 ratio, and then incubated with 40 μl HisPur Cobalt Resin for 2 hours at 4°C. The resins were then washed 4 times with wash buffer. Proteins were eluted by incubating resin in elution buffer (50 mM Sodium Phosphate pH 7.4, 300 mM Sodium Chloride, 150 mM imidazole) at RT for 5 minutes. Purified protein samples were quantified by measuring absorbance at 280 nm on a NanoDrop Spectrophotometer.

### ELISA assay for VHH binding to RBD

Maxisorp plates (BioLegend, 423501) were coated with 1μg/ml anti-Flag antibody (Sigma Aldrich, F1804) in coating buffer (BioLegend, 421701) at 4°C overnight. Plates were washed once with PBST (PBS, ThermoFisher Scientific, with 0.02% TritonX-100), a 1:1 mixture of HEK293T cell culture media containing secreted RBD-3xFlag and blocking buffer (PBST with 1% nonfat dry milk) was added to the plates and incubated at RT for 1 hour. RBD coated plates were then blocked with blocking buffer at RT for 1 hour. Plates were washed twice with wash buffer and purified VHHs-6xHis diluted in blocking buffer were added to the plates and incubated at RT for 1 hour. Plates were washed three times with wash buffer, HRP conjugated anti-His tag secondary antibody (BioLegend, 652503) diluted 1:2000 in blocking buffer was then added to the plates and incubated at RT for 1 hour. Plates were washed three times with wash buffer and TMB substrate (BD, 555214) was added to the plate and incubate at RT for 10 to 20 minutes. Stop buffer (1N Sulfuric Acid) was added to the plates once enough color developed. Quantification of plates was performed by measuring absorbance at 450 nm on a BioTek synergy H1 microplate reader. Data reported were background subtracted. Two levels of background subtraction were performed: (1) subtracting absorbance measured from wells incubated with blocking buffer only (without purified VHHs-6xHis) from sample measurements (reflecting background absorbance by plates); and (2) subtracting absorbance from each VHH incubated wells coated only with anti-Flag antibody and without RBD (reflecting non-specific binding of each VHH).

### Pseudotyped SARS-CoV-2 lentivirus production and lentivirus production for transductions

Lentivirus production was performed as previously described ^20^. Briefly, HEK293T cells were seeded at 0.8×10^6^ cells per well in a 6 well plate and were transfected the same day with TransIT®-293 Transfection Reagent and a mix of DNA containing 1 μg psPAX, 1.6 μg pTRIP-SFFV-EGFP-NLS and 0.4 μg pCMV-SARS2ΔC-gp41. Medium was changed after overnight transfection. SARS-CoV-2 S pseudotyped lentiviral particles were collected 30-34 hours post medium change and filtered on a 0.45μm syringe filter. To transduce HEK293T ACE2 the same protocol was followed, with a mix containing 1 μg psPAX, 1.6 μg pTRIP-SFFV-Hygro-2A-TMPRSS2 and 0.4 μg pCMV-VSV-G.

### SARS-CoV-2 S pseudotyped lentivirus neutralization assay

The day before the experiment, 5×10^3^ HEK293T ACE2/TMPRSS2 cells per well were seeded in 96 well plates in 100 μl. On the day of lentivirus harvest, SARS-CoV-2 S pseudotyped lentivirus was incubated with VHHs or VHH elution buffer in 96 well plates for 1 hour at RT (100 μl virus + 50 μl of VHH at appropriate dilutions). Medium was then removed from HEK293T ACE2/TMPRSS2 cells and replaced with 150 μl of the VHH + pseudotyped lentivirus solution. Wells in the outermost rows of the 96 well plate were excluded from the assay. After overnight incubation, medium was changed to 100 μl of fresh medium. Cells were harvested 40-44 hours post infection with TrypLE (Thermo Fisher), washed in medium, and fixed in FACS buffer containing 1% PFA (Electron Microscopy Sciences). Percentage GFP was quantified on a Cytoflex LX (Beckman Coulter) and data were analyzed with FlowJo.

### Affinity maturation

Error-prone PCR was used to introduce random mutations across the full length of selected VHH DNA sequences. 0.1 ng of plasmid carrying DNA sequence encoding each selected VHH were used as template in PCR reactions using Taq DNA polymerase with reaction buffer (10 mM Tris-HCl pH 8.3, 50 mM KCl, 7mM MgCl_2_, 0.5 mM MnCl_2_, 1 mM dCTP, 1 mM dTTP, 0.2 mM dATP, 0.2 mM dGTP) suitable for causing mutations in PCR products. Mutagenized library for input to CeVICA was made by ligating PCR products of error-prone PCR that carries VHH to DNA fragment containing the remaining elements required for ribosome display. Three rounds of ribosome display and *in vitro* selection were performed on the mutagenized library (pre-affinity maturation, after error-prone PCR) as described in the ***In vitro selection*** section, during which the incubation time of the binding step was kept between 5 seconds to 1 minute to impose a stringent selection condition, additional error-prone PCR was not performed during the selection cycles. The output library (post-affinity maturation) was sequenced along with the pre-affinity maturation library as described in the High throughput full-length sequencing of VHH library section.

### Identification and ranking of beneficial mutations

To identify potential beneficial mutations for each selected VHH we built an amino acid profile (a.a. profile) table for each VHH family in the pre- and post-affinity maturation library, and identified amino acids with increased frequency in the post-affinity maturation population compared to their pre-maturation frequency. For each VHH parental sequence, an a.a. profile was built of the percent of each a.a. across all VHH sequences originated from one parental VHH in the pre-affinity maturation library (“pre-a.a. profile”) and in the post-affinity maturation library (“post-a.a. profile”). A percent point change table was generated by subtracting the pre-a.a. profile from the post-a.a. profile, describing the change of frequency of each observed amino acid at each position of the VHH protein following affinity maturation.

We defined a putative beneficial mutation as either (**1**) the non-parental amino acid with the biggest increase in frequency if its increase is at least 0.5 percentage points; the score is the difference from the parental amino acid frequency; or (**2**) the non-parental amino acid with the biggest increase after the parental amino acid if the increase is at least 1.5 percentage points; the score is the percent point change of the beneficial mutation. To avoid too many proximal putative beneficial mutations (which may cause structural incompatibility), a putative beneficial mutation was discarded if it (**1**) is outside the CDRs; (**2**) is less than 3 positions away from another beneficial mutation (“nearby mutation) and has a smaller beneficial mutation score than the nearby mutation; and (**3**) co-occurs less than twice with the nearby mutation. From this final list of putative beneficial mutations, different combinations were picked and incorporated into each VHH parental sequence that include one combination of all beneficial mutations in CDRs, one combination of the top-3 ranked (by beneficial mutation score) mutations in frames, and at least one combination of both CDR mutations and frame mutations (**table S7**).

## Acknowledgements

We thank Christopher M Vockley for critical reading and editing of the manuscript, Matthew H Bakalar for helping with cloning VHH72, Leslie Gaffney and Anna Hupalowska for assistance in figure making, Michael Farzan for providing HEK293T expressing ACE2 and for discussing the SARS-CoV-2 S pseudotyped lentivirus neutralization approach, Jonathan Abraham for providing the pUC57-nCov19-S plasmid. Work was supported by the Klarman Cell Observatory and Klarman Incubator at the Broad Institute, NHGRI 5RM1HG006193 (A.R.) and HHMI (A.R.). M.G. is the recipient of an EMBO Long-Term Fellowship (ALTF 486-2018) and a Cancer Research Institute/Bristol-Myers Squibb Fellow (CRI2993). Until July 31, 2020, A.R. was an Investigator of the Howard Hughes Medical Institute.

## Author contributions

X.C. and A.R. conceived the study. X.C. designed and developed the CeVICA platform, performed selection and identification of EGFP and RBD binders, performed affinity maturation of RBD binders. M.G. developed and performed SARS-CoV-2 S pseudotyped lentiviruses neutralization assay. N.H. provided support for pseudotyped lentiviruses neutralization assay. X.C. and A.R. wrote the manuscript, with contributions from all co-authors.

## Data and materials availability

Antibody sequences are in **table S7** and will be made publicly available upon publication. Code for computational analysis will be available on Github. Key plasmids generated in this study will be deposited in Addgene.

---

**Fig. S1.**
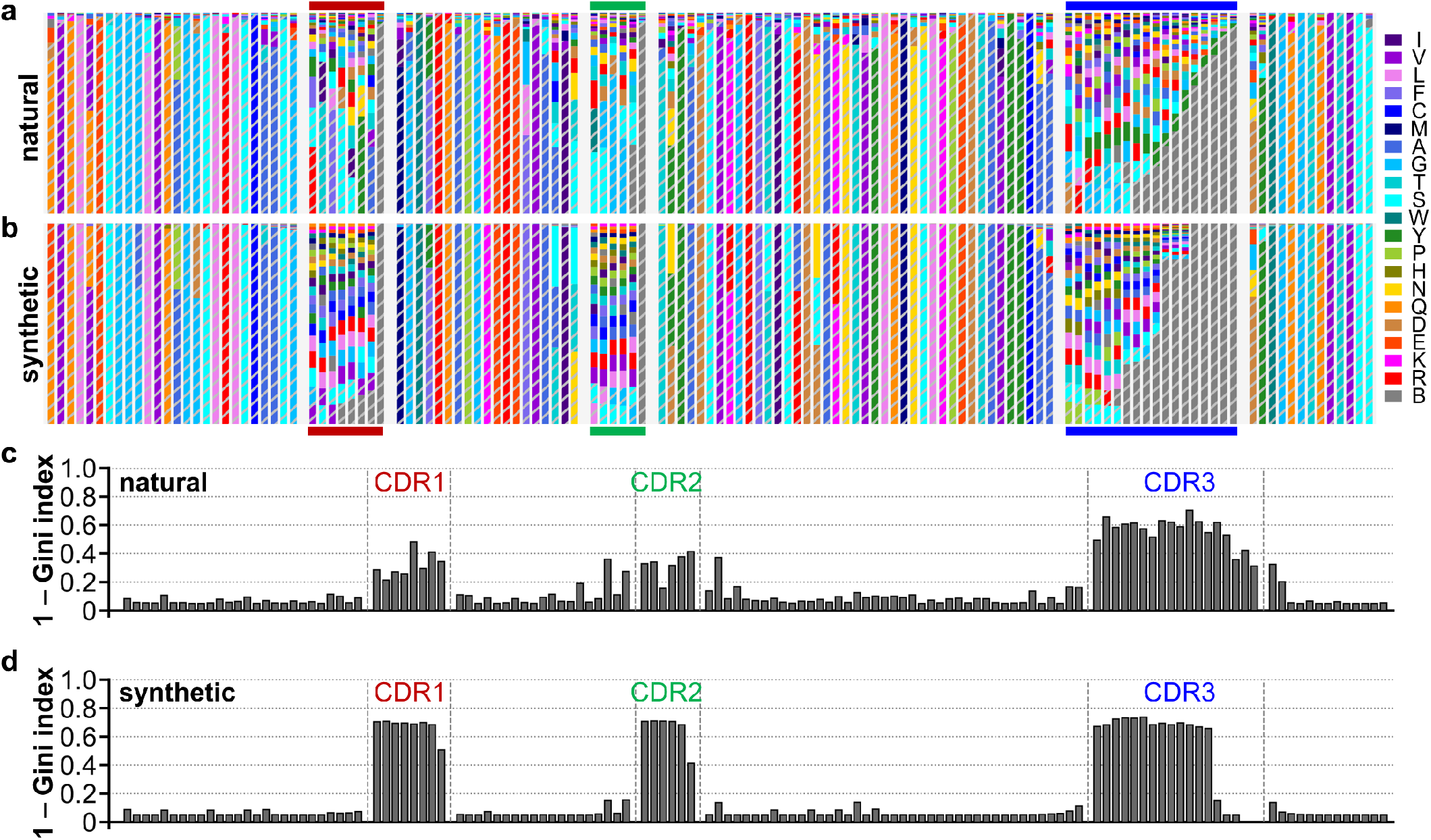
Amino acid profiles of natural and synthetic VHHs. (**a**) Position-wise amino acid profile of natural VHHs (298 VHHs, PDB) and (**b**) synthetic VHHs. Amino acids were color coded according to labels to the right, B indicates an empty position. Bar height is the relative percentage of each amino acids. The two most common amino acids were shown as patterned bars while others were shown as solid bars. (**c**) Plot of diversity index (as 1 – Gini index) for each amino acid position of natural VHHs and (**d**) synthetic VHHs.

**Fig. S2.**
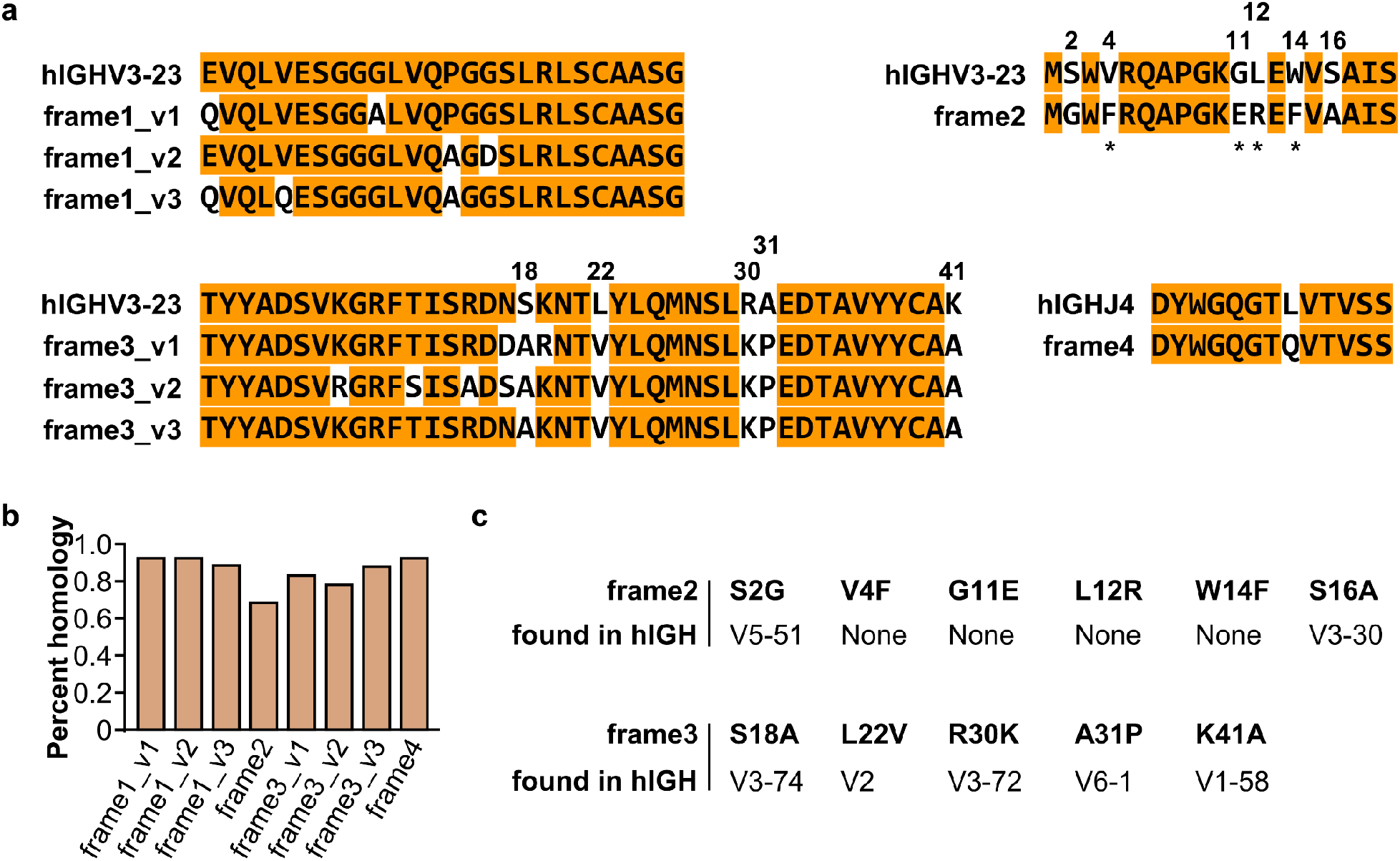
Design of VHH frames and their homology to human IGH genes. (**a**) Amino acid sequences encoded by frames that serve as templates for VHH library generation were aligned to the corresponding segments of the human IGHV3-23 (hIGHV3-23) or IGHJ4 (hIGHJ4). Positions in hIGHV3-23/hIGHJ4 that are identical to the corresponding position in at least one VHH frames are highlighted in orange. Positions in VHH frames that are identical to the corresponding position in hIGHV3-23/hIGHJ4 are highlighted in orange. hIGHV3-23 positions not identical to any VHH frames are numbered according to its position within the segment. Asterisks indicate VHH hallmark residues thought to be required for VHH’s independence of light chain. (**b**) Percent homology of VHH frames to the closest human gene. (**c**) List of VHH residues at positions numbered in (a) and representative human IGHV genes that encode the same VHH residue at the corresponding position. None: no human IGHV genes has the VHH residue at the corresponding position.

**Fig. S3.**
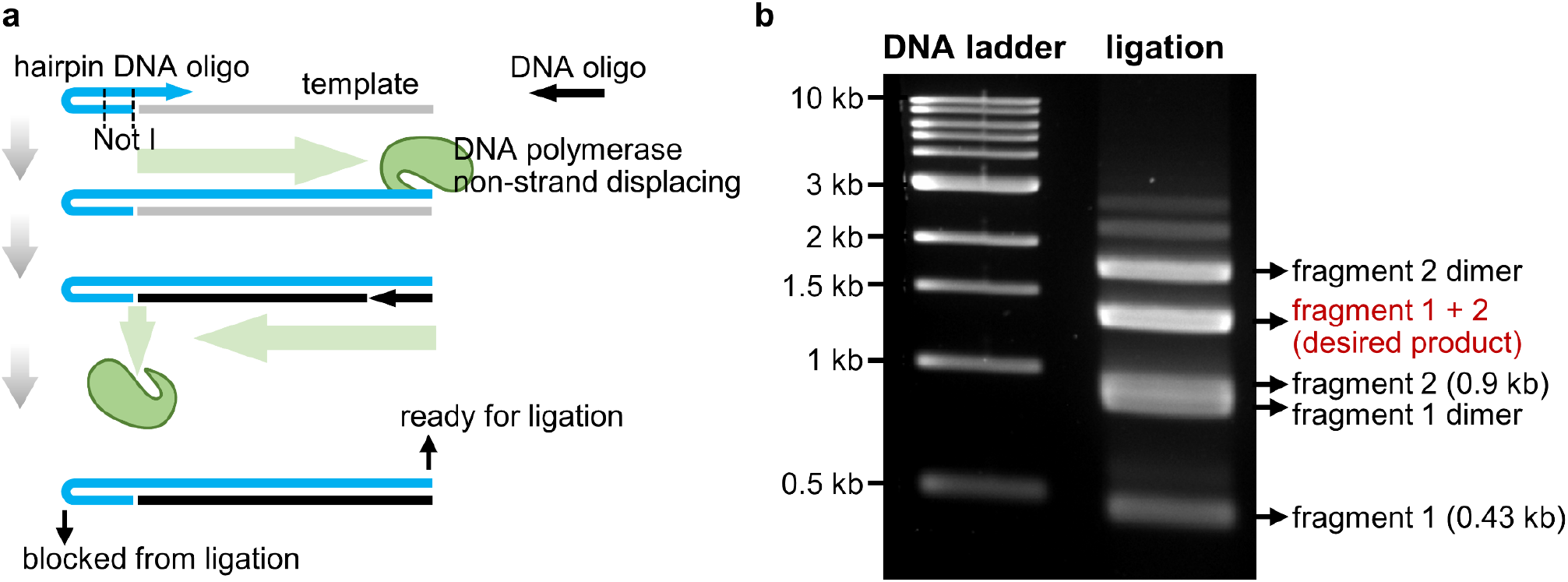
Working principles for orientation-controlled ligation by end blocking using hairpin oligos. (**a**) working principle for generating one end blocked DNA for orientation-controlled ligation by PCR using a hairpin DNA oligo. (**b**) Representative orientation-controlled ligation products visualized by agarose gel electrophoresis.

**Fig. S4.**
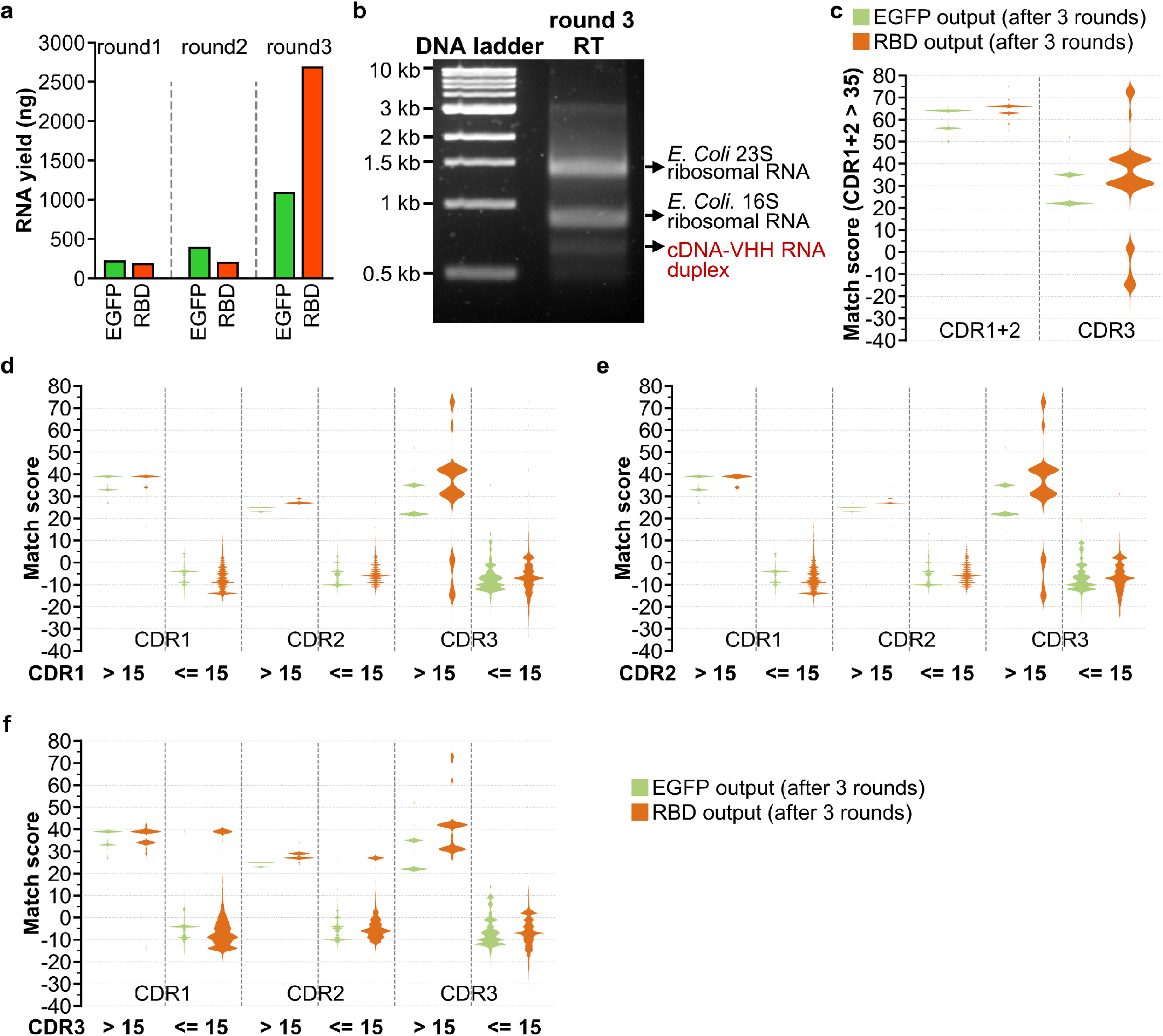
Evaluation of ribosome display and selection rounds. (**a**) Yield of recovered RNA at each round of ribosome display and selection for EGFP or RBD targets. (**b**) Representative RT reaction (without heat denaturation) product for RBD selection after 3 rounds, visualized by agarose gel electrophoresis. (**c**) Plot of match scores of sequence pairs with a combined CDR1 and CDR2 score > 35. (**d**) Plot of match scores of sequence pairs (from 2000 randomly sampled sequences) with indicated CDR1 scores, and (**e**) indicated CDR2 scores, and (**f**) indicated CDR3 scores.

**Fig. S5.**
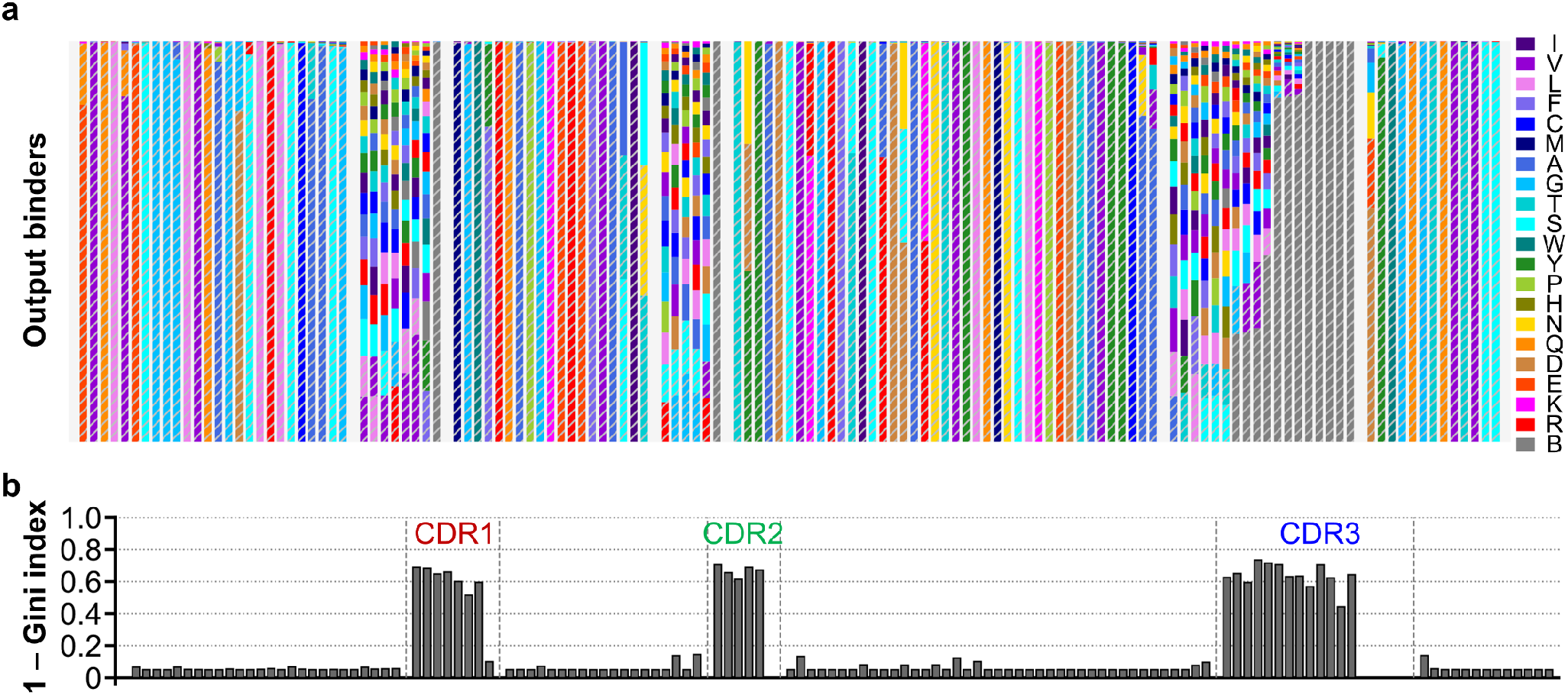
Amino acid profile for EGFP and RBD unique output binders. (**a**) Amino acid profile of representative VHH sequence for each unique cluster identified from RBD and EGFP output libraries (“output binders”, 932 sequences). Plotted as described in **Fig S1a**. (**b**) Plot of diversity index (as 1 – Gini index) for each amino acid position of output binder VHHs.

**Fig. S6.**
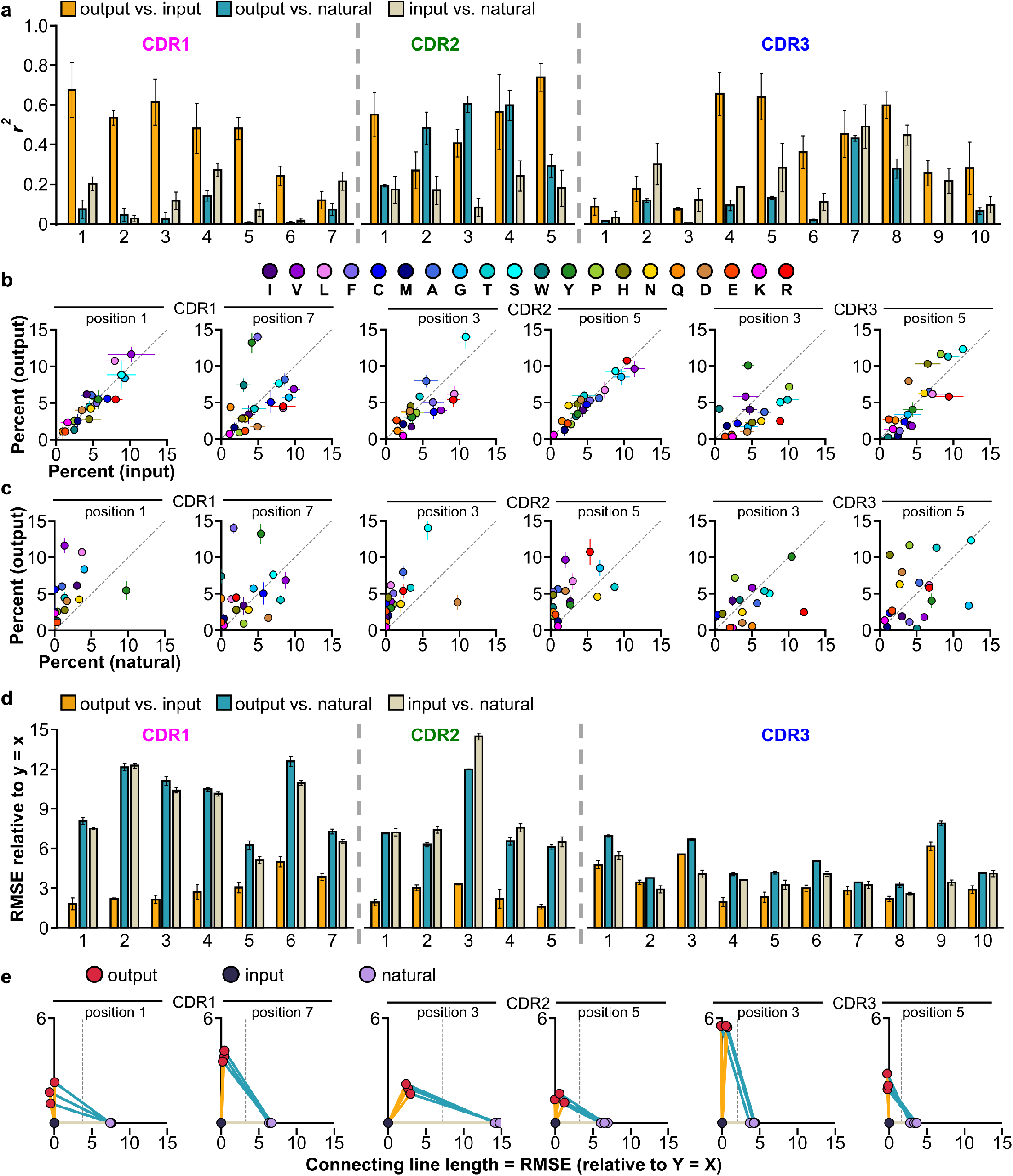
Unique output binders amino acid profiles correlate more highly with that of input library than natural VHHs. (**a**) *r*^2^ values for the amino acid percentages in the indicated sequence group pairs at each CDR position. 298 natural VHHs (natural) and 298 randomly sampled sequences from input library (input) and output binders (output) were analyzed. Three random sampling trials were performed to generate three *r*^2^ for each position. (**b**) Scatter plots of the percentage of each amino acid in the input library and the output binders and (**c**) that in the natural VHHs and the output binders at representative CDR positions. Circles are the mean and error bars are the standard deviation of data. (**d**) Root mean square error (RMSE, relative to Y = X line) values for the indicated sequence group pairs at each CDR position. Using the same randomly sampled sequences as (a). (**e**) Plot showing the similarity distances between the three sequence groups, with each connecting line length between two sequence groups indicating their RMSE. Vertical dashed lines indicate the middle point of the distance between output and natural sequence groups

**Fig. S7.**
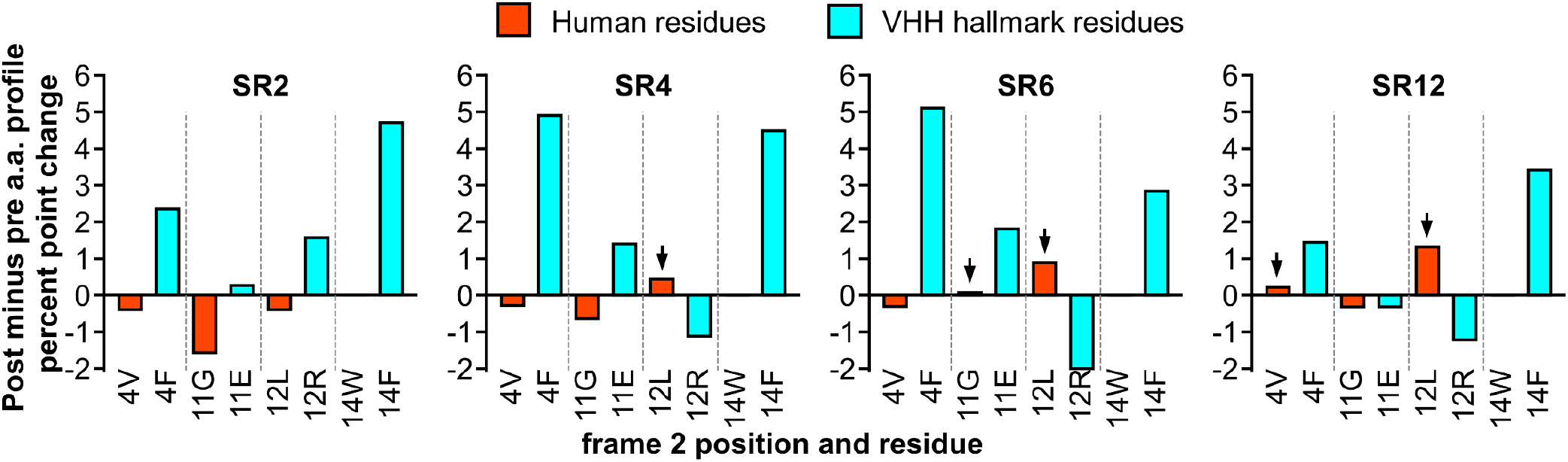
Affinity maturation leads to some VHH hallmark residues converting to the corresponding human VH residues. The post-minus pre-affinity maturation percent point change of VHH hallmark residues and the corresponding human residues for each VHH. Arrows indicate human residues with increased frequency as a result of affinity maturation.

**Table S1. Natural VHH sequences selected for calculating natural VHH amino acid profile**. Amino acid sequences of all VHHs from Protein Data Bank (sheet: all_VHH_RSCB) and from which 298 unique VHHs were selected to represent natural VHHs (sheet: unique_VHH_RSCB), the sequences were separated into 4 frames and 3 CDRs.

**Table S2. Amino acid profile of natural VHHs and synthetic VHHs in the VHH input library.** Position-wise amino acid profile of natural VHHs and VHHs in the input library. Positions are relative positions within each segment and numbers are percentage of the corresponding amino acid labelled to the left of each segment.

**Table S3. Primers and templates used for generation, selection and sequencing of VHH library.** Primer sequences used in this study and VHH frame template sequences. PCR cycling conditions were also shown.

**Table S4. List of RBD binder clusters.** A list containing key information for all predicted RBD binding clusters (sheet: all clusters) and unique RBD binding clusters (sheet: all CDR unique). Cluster ID, size, CDR representative sequences, CDR consensus sequences, CDR scores (**Materials and Methods**), and whether each CDR is unique to RBD and not found in EGFP clusters were shown.

**Table S5. List of EGFP binder clusters.** A list containing key information for all predicted EGFP binding clusters (sheet: all clusters) and unique EGFP binding clusters (sheet: all CDR unique). Cluster ID, size, CDR representative sequences, CDR consensus sequences, CDR scores (**Materials and Methods**), and whether each CDR is unique to EGFP and not found in RBD clusters were shown. The cluster with ID 0, is a spike-in VHH ^15^, and did not originate from the input library.

**Table S6. Affinity maturation subtracted amino acid profile for SR4 and SR6.** Position-wise post-minus pre-affinity maturation amino acid profile for SR4 and SR6. Numbers are percent point change of each amino acid after affinity maturation.

**Table S7. Amino acid sequences of VHH variant and the mutations they contain.** Amino acid sequences of all VHH variants characterized in this study.

**Table S8. VHH variants ELISA and neutralization data.** ELISA binding assay and pseudotyped virus neutralization assay results for all VHH variants characterized in this study.

**Table S9. High-throughput sequencing and analysis metadata.** Number of sequences obtained by high-throughput sequencing for indicated analyses.

**Data S1. Cluster files for SR1, SR2, SR4, SR6, SR8, SR12.** Text file containing all sequences belonging to each cluster. Each line in the file represent one sequence, both segments and full length of the sequence were shown, shown items were divided by “#” and in the order from start to end of each line was: CDR1 amino acid sequence, CDR2 amino acid sequence, CDR3 amino acid sequence, full-length amino acid sequence, full-length DNA sequence.

